# Hybrid untargeted and targeted RNA sequencing facilitates genotype-phenotype associations at single-cell resolution

**DOI:** 10.1101/2025.11.06.686971

**Authors:** Jiayi Wang, Maria Constanza Maldifassi, Anna Bratus-Neuenschwander, Qin Zhang, Felix Beuschlein, David Penton, Mark D. Robinson

## Abstract

Long-read scRNA-seq enables simultaneous, unbiased identification of transcriptomic variants and gene expression but is often limited by low read coverage, restricting genotype–phenotype analyses at single-cell resolution. We systematically evaluated short-read whole-transcriptome amplification (SR-WTA) via Illumina, long-read whole-transcriptome amplification (LR-WTA), and long-read targeted sequencing (LR-Twist) via PacBio for cell typing and genotyping performance. Based on these comparisons, we propose a hybrid strategy and accompanying Snakemake pipeline that integrates SR-WTA with LR-Twist to leverage the strengths of both approaches. SR-WTA provides broad transcriptome coverage, while LR-Twist enriches a 50-gene panel for deeper variant detection. This hybrid approach improves power to link mutational profiles with transcriptional programs at single-cell resolution.

## Background

Single-cell sequencing technologies have transformed our ability to characterize cellular identity and function^[1][2]^. Among these, single-cell RNA sequencing (scRNA-seq) is widely used to capture transcriptional variability and cellular heterogeneity, and it also provides some (albeit limited) opportunity for variant detection. Compared with scDNA sequencing, scRNA-seq is more cost-effective and enables the simultaneous assessment of genetic and transcriptomic variants in the same cells. However, beyond the limitations of genotyping from RNA, short-read scRNA-seq is rather limited for variant detection because library fragmentation introduces 5’ and 3’ biases, restricting the reach to link mutation status with expression profiles. This link is critical for understanding how genetic variation shapes cellular phenotypes and tumor heterogeneity^[3,4]^.

Long-read scRNA-seq overcomes these capture biases by sequencing full-length transcripts, improving the ability to detect variants. Yet, it generally suffers from lower sequencing depth^[5][6]^, which can limit variant calling in lowly expressed genes and constrain the number of cells genotyped. Early targeted RNA sequencing approaches using hybridization capture enabled enrichment of specific transcripts for detailed characterization^[7][8][9]^. Single-cell approaches typically barcode cDNA using established methods such as 10x Genomics or Drop-seq^[10][11]^, allowing integration across sequencing technologies. Building on this, several studies^[12][13][14][15][16]^ have shown combined targeted long-read scRNA-seq with whole-transcriptome short-read scRNA-seq to jointly characterize cell-type diversity and mutation-associated expression profiles. To date, most targeted approaches rely on Oxford Nanopore Technologies (ONT) platform, which, despite lower cost, has higher sequencing error rates compared to Pacific Biosciences (PacBio)^[17]^, thus leading to lower detection of single nucleotide polymorphisms (SNPs) and insertion or deletion events (indels)^[18][19]^.

We present a proof-of-principle comparison of three scRNA-seq strategies: Illumina short-read whole-transcriptome sequencing (SR-WTA), PacBio Kinnex long-read whole-transcriptome sequencing (LR-WTA), and Twist hybridization-based targeted capture of 50 genes relevant to steroidogenesis applied to PacBio Kinnex libraries (LR-Twist). Using the same cDNA generated with the 10x Genomics single-cell 3’ kit, we directly compared the performance for cell type annotation and detection of small variants (Fig. 1a). Our goal is to assess their complementarity and propose a practical strategy and computational pipeline for single-cell genotype-phenotype integration.

**Fig. 1:**
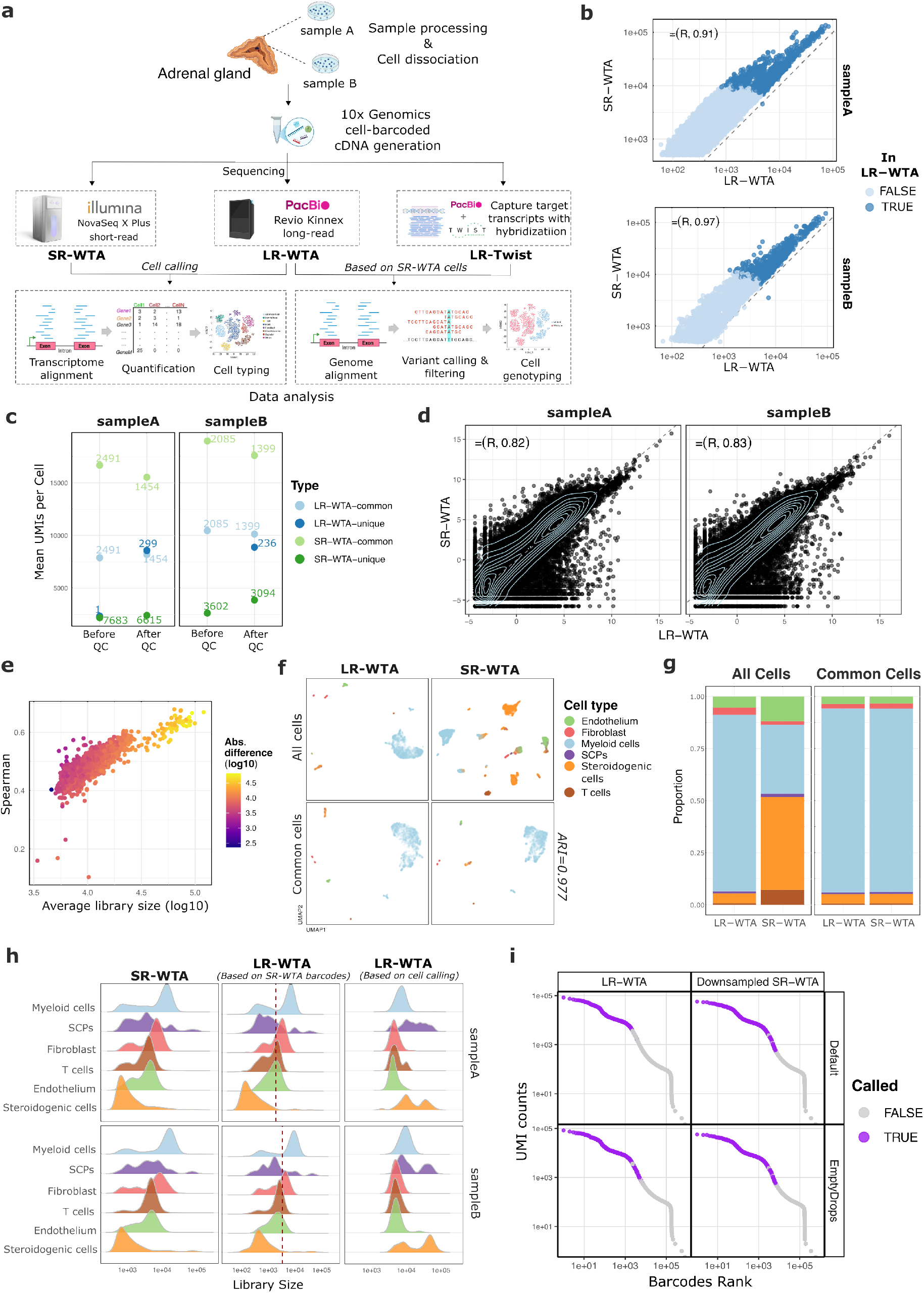
Comparison between SR- and LR-WTA. (**a**) Schematic workflow of the three scRNA-seq technologies and data analysis. (**b**) Comparison of per-cell library sizes (summed UMIs) between SR-WTA and LR-WTA, based on SR-WTA cells, with coloring indicating whether cells were also detected in LR-WTA. (**c**) Mean UMI counts per cell for common and unique cells in SR-WTA and LR-WTA, with labels indicating the number of cells, before and after QC. (**d**) Scatter plot comparing gene-level expression (logCPM) between SR-WTA and LR-WTA, with Spearman correlation coefficient indicated. (**e**) Relationship between average library size (SR-vs. LR-WTA) and cell–cell Spearman correlation on gene expression, colored by the absolute difference in library size between the two. (**f**) UMAP projection of cells from samples A and B, shown either for all cells (based on their respective cell calling) or for the common cells between LR-WTA and SR-WTA. The projection is based on the top 3,000 highly variable genes (HVGs), with cells colored by cell types annotated using *CellTypist*. (**g**) Cell type proportions in LR-WTA and SR-WTA, stratified by all vs. common cells. (**h**) Scaled distribution of UMI counts per cell (library size) stratified by cell type in SR- and LR-WTA (based on SR-WTA or LR-WTA cell calling) for samples A and B. The dashed line indicates the minimum library size after cell calling and QC in LR-WTA, i.e. minimum library size in the third column. (**i**) Cell barcode rank plot for sample A from LR-WTA and downsampled SR-WTA data (Seed 1). Cells are colored according to whether they were called by the default cell-calling methods (*CellRanger* for SR-WTA or *isoseq* for LR-WTA) and by the same *EmptyDrops* method. Panel **f, g** and **h** use the same color legend.

## Result and Discussion

### Overview of the workflow

Briefly, we utilized shared single-cell cDNA libraries derived from a single adrenal gland to compare the performance of SR-WTA, LR-WTA, and LR-Twist for different analytical goals. Reads were aligned to both the transcriptome and the genome, linking each sequencing strategy to its analytical purpose; transcriptome alignment enables quantification and cell type annotation, while genome alignment supports variant detection (at the pseudobulk level). To evaluate the genotyping performance, we selected candidate variants based on read coverage, quality, coding-region restriction, and absence from common variant databases, allowing a direct comparison of variant calls from targeted capture versus whole-transcriptome sequencing. We compared cell-typing results obtained with SR-WTA and LR-WTA, which capture whole-transcriptomic profiles, and cell-genotyping results between LR-WTA and LR-Twist, which allow detection of variants across full-length transcripts (Fig. 1a, Fig. S1).

### SR-WTA vs. LR-WTA

We first assessed the sequencing quality of SR- and LR-WTA. Standard processing workflows were applied, with SR-WTA and LR-WTA processed using *CellRanger* and *isoseq*, respectively. Both workflows implement similar steps but in a different order (Table S1). The SR-WTA library generated ∼242.8 million reads per sample, whereas the LR-WTA library produced ∼81.7 million segmented reads per sample (from HiFi read deconcatenation; see Methods). Although LR-WTA yielded fewer reads than SR-WTA, the median read length of LR-WTA (∼853 bp) is 2.8 times longer than that of SR-WTA (300 bp, pair-end), resulting in a comparable total number of sequenced bases. Before alignment, long reads were preprocessed and deduplicated via clustering on cell barcodes and unique molecular identifiers (UMIs). A similar deduplication step was performed after alignment for SR-WTA. Sequencing saturation, defined as 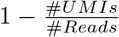, was on average 13% and 36.4% for LR- and SR-WTA, respectively. Lower sequencing saturation indicates that additional sequencing would recover more transcripts. In addition to deduplication, long-read datasets undergo cell calling prior to alignment, which further reduced the number of reads to an average of ∼27.6 million for LR-WTA (Fig. S2). For SR-WTA, by contrast, cell calling is performed after alignment. The alignment rate for LR-WTA reads to GRCh38.p14 genome remained high, averaging 99.6%, compared to 95% for SR-WTA reads (Table S2, Table S3).

Differences in sequencing depth, preprocessing order, and the inherent characteristics of cell-calling methods led to variations in the number of cells identified (via cell barcodes) in each dataset. Although we observed a very high correlation between the library size between SR- and LR-WTA (Spearman=∼0.94), SR-WTA yielded substantially more cells, with additional 7683 and 3602 cells in sample A and sample B, respectively, compared to LR-WTA (Fig. 1b, Fig. 1c). Both *CellRanger* and *isoseq* use the knee-finding method by default. However, after setting the threshold based on the barcode rank plot, *CellRanger* applies an additional RNA-profile-based filter that reselects barcodes falling below the threshold if their transcriptomic profiles differ significantly from ambient RNA, thereby recovering more real cells^[20][21]^. Consequently, we noted that SR-WTA-unique cells exhibited lower mean UMI counts than cells that were also detected in LR-WTA. Moreover, on average, 260 common cells per sample passed quality control (QC; see Methods) in LR-WTA but not in SR-WTA (Fig. 1c). Thus, SR-WTA recovers more cells compared to LR-WTA, though these may be of relatively lower depth.

We next evaluated the gene-by-cell count matrices constructed from the SR- and LR-WTA datasets to assess their potential for annotating cells. We first compared gene-level expression profiles using logcounts per million (logCPM). As previously noted, SR-WTA recovered more cells overall; however, many of these were of low depth. Despite this difference, both platforms exhibited substantial agreement in pseudobulk gene expression, both when aggregating across all cells and when assessed within annotated cell types (see below). While SR-WTA exhibited higher expression for most genes, its overall expression levels were highly consistent with LR-WTA, showing a strong linear relationship with an average Spearman’s correlation coefficient of 0.825 (Fig. 1d, Fig. S3). After QC, 11,043 genes and 3,388 cells were retained for LR-WTA, and 13,542 genes and 12,562 cells for SR-WTA, with 9,961 genes and 2,852 cells shared between the two platforms. Among these cells, we compared gene expression by correlating cell-wise log-normalized counts profiled with SR-WTA and LR-WTA, considering the union of genes detected. Cell-wise Spearman correlations between SR- and LR-WTA were generally high (mean = 0.5), indicating moderate agreement. Correlation strength was influenced by both the average library size and the difference in library size between the two platforms, with cells having lower average library sizes or larger library size discrepancies tending to exhibit weaker correlations (Fig. 1e). We then projected gene expression profiles into two-dimensional space using UMAP, using either all cells/features or only those shared between the two datasets (Fig. 1f). SR-WTA shows more distinct subpopulations than LR-WTA, likely due to deeper transcriptional coverage. To quantify the similarity in low-dimensional space, we computed pairwise Euclidean distances from the first 20 principal components and correlated the corresponding distances between SR-WTA and LR-WTA. This yielded a Spearman’s correlation of 0.97, indicating strong agreement. Together, these results indicate that SR- and LR-WTA generated similar expression profiles, allowing for comparable cell type classification.

Lacking ground truth, we compared cell type annotation obtained from *CellTypist* ^[22][23]^ using the human adrenal gland reference model^[24]^. Of the nine reference cell types, six were identified in both SR-WTA and LR-WTA datasets. In LR-WTA, most cells were annotated as myeloid, whereas steroidogenic cells predominated in SR-WTA. For cells shared between SR-WTA and LR-WTA, annotations were highly consistent, with an adjusted rand index (ARI) of 0.977 and similar proportions across cell types (Fig. 1f, Fig. 1g). Those additional cells detected in SR-WTA were mostly steroidogenic, consistent with their lower RNA content (Fig. 1h). These cells were likely recovered in SR-WTA due to deeper sequencing and more sensitive cell calling in *CellRanger*, whereas they were undersampled in LR-WTA.

The higher number of detected cells in SR-WTA appears to be driven by both sequencing depth and differences in cell-calling strategies. To investigate this, we downsampled SR-WTA to match the LR-WTA sequencing depth on a per-sample basis, repeating the procedure five times to account for sampling variability. Using the default cell-calling pipelines (*isoseq* for LR-WTA and *CellRanger* for SR-WTA), downsampled SR-WTA recovered less cells but still more than LR-WTA. This difference is largely attributable to *CellRanger* calling additional cells below the knee threshold (Fig. 1i, Fig. S4a,b). Annotation of these cells indicates that they are predominantly low–RNA-content steroidogenic cells (Fig. S4c,d). We then applied the same cell-calling method (*EmptyDrops*) to both datasets. Under this unified approach, 10583 and ∼10779 cells were called for LR-WTA and downsampled SR-WTA datasets, with ∼77% and ∼84% overlap for sample A and B, respectively (Fig. 1i, Fig. S5a,b). Cell-type separation and proportions were also more consistent (Fig. S5c,d). The remaining non-overlapping cells likely arise from sampling variability during downsampling, alignment differences, and technology-specific processing orders.

Overall, SR-WTA shows strong consistency with LR-WTA in transcriptional profiles and cell type annotation, while recovering substantially more cells, primarily due to deeper sequencing depth and differences in cell calling strategies.

### LR-WTA vs. LR-Twist

We next investigated how effective Twist-based targeted sequencing captured the intended genes. As expected, the total number of raw reads in the LR-Twist libraries was considerably lower, with approximately 31.85 million segmented reads, compared to 81.7 million in the LR-WTA datasets. WTA libraries are typically sequenced at greater depth since they capture the entire transcriptome, whereas targeted sequencing focuses only on the interested regions. To minimize differences from independent cell calling while maximizing cell recovery, we retained genome-aligned LR-WTA and LR-Twist reads whose cell barcodes matched those identified in SR-WTA. In LR-Twist, 38.5% and 27.9% of UMIs mapped to on-target genes in samples A and B, respectively, compared with only 0.5% and 0.4% in LR-WTA (Fig. 2a), representing an approximately 70-fold enrichment. Because the targeted genes are associated with steroidogenic cells, this cell type showed the strongest enrichment, as expected (Fig. S6). This enrichment translated into roughly an eightfold increase in UMI counts for target genes on average, observed at both the gene and transcript levels (Fig. S7, Fig. S8). Nonetheless, off-target genes were also captured in proportion to their native expression (Fig. 2b). Similar observations have been reported in other targeted sequencing approaches^[13][25]^. Overall, these results demonstrate Twist-based targeted sequencing achieves efficient transcript capture, even with a panel of 50 genes.

**Fig. 2:**
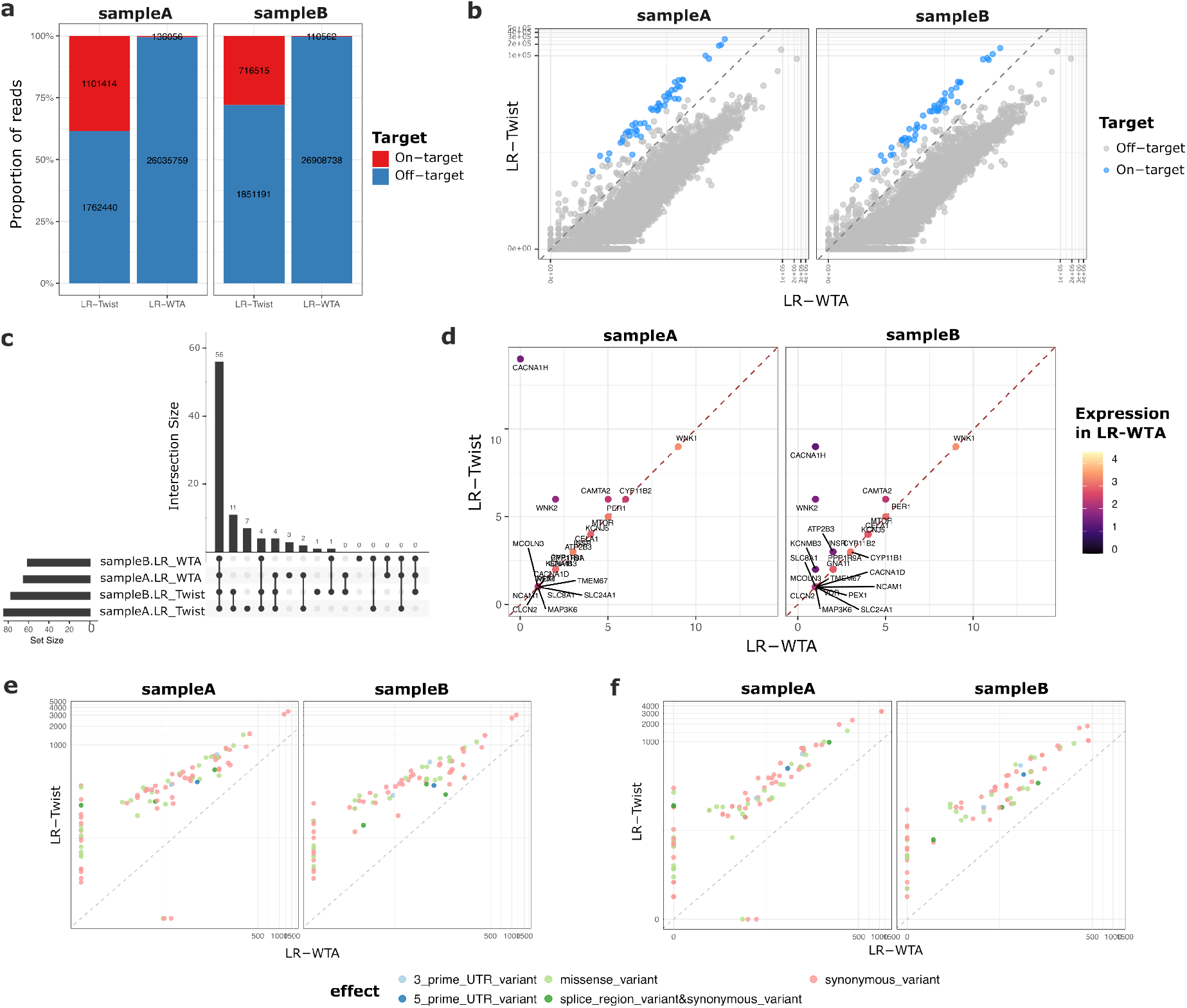
Comparison between LR-WTA and LR-Twist. (**a**) Proportion of UMI counts mapping to on- and off-target genes in LR-Twist and LR-WTA, with labels indicating the number of UMIs. (**b**) Scatter plot comparing summed gene-level UMI counts across all cells between LR-WTA and LR-Twist (on log10 scale), colored by on- and off-target genes. (**c**) UpSet plot comparing the number of variants identified on each sample using LR-WTA or LR-Twist. (**d**) The number of variants identified on each targeted genes, colored by the gene expression in LR-WTA. (**e**) Number of reads mapped and (**f**) number of cells can be genotyped on each variant locus, colored by annotated variant effect.

Finally, we compared the performance of LR-WTA and LR-Twist to decipher genotype information based on variant count, read depth, and number of genotyped cells. For demonstration, we focused on candidate variants filtered by read depth, quality, and coding regions. In total, there were 21 variants only identified in the LR-Twist samples, with 7 and 3 of them only identified in sample A and B, respectively (Fig. 2c). As noted, variant calling from RNA-seq is constrained by gene expression levels and possibly allele specificity (e.g. a region may be heterozygous but only one expressed allele is observed). Lowly expressed genes generate insufficient reads, reducing detection sensitivity. Accordingly, one advantage of LR-Twist is that it enables detection of variants in such lowly expressed genes. For example, no variants were detected in *CACNA1H* due to its low expression with total 3 UMIs in the LR-WTA data. With LR-Twist, we detected 9 and 15 variants in CACNA1H in sample A and B, respectively (Fig. 2d). However, we also observed variants identified in LR-WTA but not LR-Twist. One such case is the SNV in *CYP11B2* (p.His439Tyr) in sample A. In LR-WTA, this site had 6 reads supporting the alternative allele (1 forward, 5 reverse) and 16 reads supporting the reference allele (1 forward, 15 reverse). LR-Twist provided much deeper coverage, with 48 reads supporting the alternative allele (5 forward, 43 reverse) and 308 reads supporting the reference allele (8 forward, 300 reverse). The forward/reverse distribution indicates a strand imbalance, with Fisher strand scores^[26]^ of 3.18 (*p* = 0.48) for LR-WTA and 16.95 (*p* = 0.02) for LR-Twist. Increased coverage in LR-Twist likely accentuated this imbalance, yielding a higher Fisher score(LR-WTA ∼27% vs. LR-Twist ∼13%; details see Supplementary 1.2) and a lower observed variant allele frequency, which together may have prevented a confident variant call in LR-Twist. Given that this variant has been documented previously^[27]^, we cannot exclude the possibility of false negatives in LR-Twist, although LR-Twist generally improves detection of known variants. To guard against both false positives and false negatives at such loci, we recommend manual inspection in Integrative Genomics Viewer (IGV) before drawing conclusions.

Notably, for many genes, we identified the same number of candidate variants in LR-WTA and LR-Twist (Fig. 2d). This is because we required only a minimum read depth of five to call a variant at the pseudobulk level. Although LR-Twist did not substantially increase the number of candidate variants in well-expressed genes, it markedly improved variant read depth. This improvement directly affected the number of cells that could be genotyped (Fig. 2e). A cell was classified as mutated for a given variant if at least one read carried the alternative allele, and as reference if only reads with the reference allele were observed. Consequently, LR-Twist increased the number of genotyped cells, thereby raising the proportion of steroidogenic cells successfully genotyped (Fig. 2f, Fig. S9). The deeper coverage not only increased sensitivity at lowly-expressed regions, as shown above, but also enabled the detection of more mutated cells, thereby strengthening the possibility to link genotype and phenotype and facilitating downstream (differential expression) statistical analyses.

## Discussion

Taken together, LR-Twist efficiently captures targeted genes and improves genotyping yield. However, on average, 65% of reads originated from off-target regions. This can be attributed to several factors: off-target transcripts may contain sequences similar to targeted regions and bind capture oligos or primers; genes located near targeted loci may be co-captured; and additional off-target reads may arise from sequencing background^[13][25]^. Other targeted sequencing approaches, such as TEQUILA-seq^[28]^ and Agilent-based enrichment^[29]^, have reported on-target rates exceeding 80%, which are higher than those observed for LR-Twist in our study. However, direct comparison is challenging due to differences in panel design, target composition, and experimental context. For example, TEQUILA-seq profiled 468 actionable cancer genes, and capture efficiency also varies depending on the specific gene set targeted. In addition, enrichment performance may differ between bulk and single-cell applications due to differences in input material, library complexity, and experimental design. The present data do not allow us to distinguish whether certain gene sets are inherently more difficult to capture or whether methodological differences between platforms account for the observed discrepancy. While our 50 gene panel achieves deep sequencing saturation, ideal for detecting rare variants, it introduces a distinct technical profile compared to broader exome or large gene panels. The “off-target paradox” inherent in low-complexity bait libraries can lead to higher relative non-specific binding, as fewer probes must compete against a vast pool of non-target cDNA. Furthermore, the high per-gene coverage and extensive PCR cycling required for small panels can amplify library preparation artifacts, such as 3’-biased RT truncations and chimeric molecules, which may be mistaken for novel isoforms. Despite these challenges, the high-resolution data provided by our targeted approach offers unparalleled granularity for our specific genes of interest, provided that rigorous bioinformatic filtering is applied to distinguish stochastic noise from true biological heterogeneity.

Given the observed correlation between LR-WTA and LR-Twist in off-target gene expression, one might consider using LR-Twist alone for both genotyping and cell typing. However, in practice, the targeted enrichment skews expression profiles and misleads cell typing (Fig. S10). As mentioned above, we also did not pursue genotyping from SR-WTA because reads are concentrated at the 3’ end, limiting variant calling outside the tagged region (Fig. S11). Therefore, we recommend a hybrid strategy in which SR-WTA provides comprehensive cell typing and LR-Twist for unbiased and targeted genotyping, together enabling robust linkage of genotype and phenotype on the single-cell resolution.

## Conclusion

In this study, we evaluated different sequencing technologies for simultaneous cell typing and genotyping, and we propose a promising strategy for linking single-cell genotypic and transcriptomic states by combining SR-WTA and LR-Twist. The workflow assumes a predefined targeted gene panel as input for capture and analysis, which in our case came from expert knowledge and literature searches. We selected genes of interest for the capture panel based on their known association with steroidogenesis. The same framework can be readily extended to other applications by targeting genes relevant to specific biological processes or disease-associated mutation hotspots. Alongside the experimental strategy, we provide a modular Snakemake pipeline that takes Illumina FASTQ files and PacBio BAM files as input, together with standard reference resources (e.g., reference genome and gene annotation), and generates expression count matrices and identifies cells carrying specific variants (Fig. S1). By providing both broader and deeper coverage of the transcriptome, SR-WTA enables the detection of additional genes and cell types, with particular advantages in recovering low-RNA-content cell types that are poorly represented in LR-WTA. LR-Twist efficiently captured and enriched a targeted panel of 50 genes, including lowly expressed ones such as *CACNA1H*. The increased coverage of targeted transcripts mitigates the risk of missing variants from lowly expressed genes and enables the detection of more mutated cells. The latter is particularly important for investigating the effects of mutations on expression programs, as it increases the statistical power when comparing mutated and reference cells; that said, the identification of mutated cells may still be challenged by allele-specific expression of the reference allele, where it comes down to sampling. In addition, LR-Twist offers a cost advantage by requiring fewer sequencing reads, as only targeted genes are captured. This reduction in sequencing depth enables multiplexing of several samples on a single PacBio SMRT Cell, in contrast to LR-WTA, which generally requires one SMRT Cell per sample.

Whereas previous studies primarily compared targeted sequencing with SR-WTA, our work extends this by incorporating LR-WTA, thereby providing a more comprehensive evaluation that considers both short- and long-read whole-transcriptome strategies. In addition, we focus on a broader target panel and on variant detection, rather than limiting the analysis to a few genes with known mutations. This demonstrates the potential of our workflow for identifying novel variants. We anticipate that focusing on an even smaller number of targets would further improve enrichment and sensitivity. This workflow may be broadly useful in genomic applications where coverage and read length are limiting factors. For instance, isoform discovery could benefit from improved detection of alternative transcripts, and structural variant analysis could also be facilitated.

## Methods

### Sample preparation

Two samples from one adrenal gland were obtained following adrenalectomy at the University Hospital Zurich. After surgical removal, samples were collected and immediately transferred to the laboratory on ice using *MACS Tissue Storage Solution* (Miltenyi Biotec, Cat# 130-100-008). Tissues were processed within 24 h of arrival in the laboratory, and were stored at 4 °C.

For sample dissection, tissues were cleaned of surrounding adipose tissue, blood vessels, and connective tissue using scalpels and forceps. *DMEM/F12* (Gibco, Cat# 11320-074) was used to keep tissues moist throughout dissection. Tissues were then minced into fragments smaller than 0.5 mm.

For the enzymatic digestion, minced tissues were digested in a solution of 1 mg/mL Collagenase I (Millipore, Cat# SCR103) prepared in a 1:1 mixture of *DMEM/F12* without calcium, with phenol red, no serum (Gibco, Cat# 21068-028), and *DMEM/F12* with serum, without phenol red (Gibco, Cat# 11039-021). Samples were incubated at 37 °C for 1 h in a water bath, with manual agitation every 15 min. Following digestion, tissues were further dissociated manually using a sterile plastic Pasteur pipette. Residual fat or visible blood vessels were removed. Cell suspensions were filtered through *MACS SmartStrainers* (70 *µ*m; Miltenyi Biotec) and washed with 25 mL ice-cold complete *DMEM/F12*. Cell suspensions were then centrifuged at 300 × g for 7 min at 4 °C. Pellets were resuspended in 25 mL of ice-cold *DMEM/F12* and centrifuged again under the same conditions. Pellets were resuspended in 2 mL of ice-cold *DMEM/F12* and incubated with 1× RBC lysis buffer (ThermoFisher, Cat# 00-4333-57) on ice for 7 min. Lysis was quenched by adding 25 mL of ice-cold PBS. Samples were centrifuged at 600 × g for 10 min at 4 °C, and pellets were resuspended in 1.5 mL of 0.05% BSA in PBS. Cell concentration and viability were assessed using trypan blue exclusion. Prior to sorting, cells were stained with DAPI (1 *µ*g/mL) to exclude dead cells. A negative control tube was always included, and only viable cells were sorted.

### GEMS-RT-cDNA obtention

Following sample cell obtention, the *Chromium GEM-X Single Cell 3’ Reagent Kits v4* (10x Genomics, USA) was used to generate single cells with barcoded gel beads in emulsion droplets (GEMs) according to the manufacturer’s protocol, allowing for reverse transcription and the generation of uniquely indexed cDNA libraries from each cell simultaneously. Cell viability and concentration of the single-cell suspensions were determined using a Luna FX7 automated cell counter with acridine orange/propidium iodide (AO/PI) staining. All samples showed ≥80% viability. Cell concentration was adjusted to achieve the desired recovery of approximately 6,000 cells per sample. Cells, reverse transcription (RT master mix) reagents, and barcoded gel beads were loaded onto a *Chromium Single Cell Chip* and processed on the *Chromium Controller X* to generate Gel Beads-in-Emulsion (GEMs), each containing a single cell, a uniquely barcoded bead, and lysis/RT reagents. Within each GEM, cell lysis and reverse transcription occurred using a thermal cycler, incorporating a cell-specific barcode and a unique molecular identifier (UMI) into each cDNA molecule. After reverse transcription, the emulsion was broken using the provided recovery agent, followed by cleanup with *Dynabeads MyOne SILANE* (included in the kit) and elution in the specified elution solution. The resulting cDNA was amplified using the specified cDNA Amplification Reaction Mix and incubated in the thermal cycler as per the kit specifications using 12 PCR cycles, then purified using *SPRIselect* beads (Beckman Coulter, Cat# B23317) together with magnetic separators. Finally, samples were run on a High Sensitivity D5000 ScreenTape using the Agilent 4200 TapeStation (Agilent, USA) for cDNA quality control and quantification.

### Library Preparation and Sequencing

#### SR-WTA

The 10x full-length cDNA was used to produce dual index short read sequencing libraries according to the manufacturer’s protocol with Chromium GEM-X Single Cell 3^*′*^ Reagent Kits v4 (10x Genomics, USA). Briefly, the cDNA was enzymatically sheared to a target size of 200–300 bp, followed by end repair and A-tailing, adapter ligation, a sample index polymerase chain reaction (PCR), and SPRI bead clean-ups with double-sided size selection. The sample index PCR added a unique dual index for sample multiplexing during sequencing. The final libraries contained P5 and P7 primers and were used in Illumina bridge amplification. Sequencing was performed using paired-end 150 bp sequencing on an Illumina NovaSeq X Plus sequencer to achieve ∼ 50,000 reads per cell.

#### LR-WTA

Single cell full-length cDNAs from two samples (sample A and sample B) with the amount of 15ng/sample (produced following 10x Chromium single cell 3’ cDNA worflow (as discribed above)), were further used to generate PacBio Kinnex single cell RNA libraries with PacBio Kinnex single-cell RNA kit (Pacific Biosciences, CA, USA), following the manufacture’s instructions from ‘Procedure & checklist — Preparing Kinnex libraries using Kinnex single-cell RNA kit’. In summery, template-switch oligo (TSO) priming artifacts generated during 10x Genomics cDNA synthesis were removed in the PCR step with a modified PCR primer (Kinnex 3’ capture primer) to incorporate a biotin tag into desired cDNA products followed by their capture with streptavidin-coated Kinnex beads. cDNA free of TSO artifacts (with the amount of 25ng/sample) was further used to incorporate the programmable segmentation adapter sequences (Kinnex primers) in 16 parallel PCR reactions for each sample, followed by directional assembly of amplified cDNA segments into a linear array. Such formed Kinnex arrays with an average size of 15 kb were further DNA damage repaired and nuclease treated to produce final Kinnex single cell RNA libraries, the quantity and quality of which were measured by the Qubit 1X dsDNA High Sensitivity Kit (Thermo Fisher Scientific, USA) and the Femto Pulse pulse-field capillary gel electrophoresis system (Agilent, USA), respectively. Each Kinnex single cell RNA library was used to prepare the DNA-polymerase complex using Revio binding chemistry v.13 (Pacific Biosciences) and further sequenced on a single 25M SMRT cell (Pacific Biosciences), on the Revio sequencer (Pacific Biosciences) yielding 5.4 M HiFi reads/ 85.2 M segmented reads and 5 M HiFi reads/ 78.2 M segmented reads for sample A and sample B, respectively.

#### LR-Twist

PacBio long-read single-cell RNA libraries were generated using a Twist custom capture panel targeting 50 genes: *ATP1A1, ATP2B3, CACNA1D, CACNA1H, KCNJ5, GNA11, MCOLN3, CLCN2, CTNNB1, CYP11B2, FRRS1L, CELA1, UBTD1, NCAM1, CAMTA2, TMEM67, KCNA4, KCNH3, KCNK5, PER1, WNK2, KCNMB3, INSR, PDE8A, PPP1R9A, USP3, NR0B2, AQP11, NFATC4, STAR, CYP11B1, KCNK3, SLC8A1, CALM1, CREB1, PRKACA, MTOR, GNAQ, SLC24A1, PEX1, VDR, TSC2, CDKN2B, MAP3K6, UBE2Q2L, SLC30A1, NR5A1, HSD3B2, NR4A2, WNK1*. Libraries were prepared using the Twist Exome Enrichment Kit (Twist Bioscience, USA) following a modified version of the protocol described in the PacBio Technical Note, “Preparing Kinnex single-cell libraries with Twist Exome Enrichment Kit” (available at: https://www.pacb.com/wp-content/uploads/Technical-note-Twist-exome-for-single-nuclei-RNA-seq-using-Kinnex.pdf). Target genes were chosen according to their relevance to steroidogenic pathways, incorporating both recognized key regulators and promising novel candidates supported by expert insight and published evidence. Breifly, full-length single cell cDNA, free of TSO artifacts (produced as discribed above) from sample A and sample B with the amount of ∼90 ng/sample was further used in pre-capture PCR with 10x cDNA Primer FWD (CTACACGACGCTCTTCCGATCT) and 10x cDNA Primer REV (AAGCAGTGGTATCAACGCAGAG) in 6 cycles reaction in order to increase the cDNA mass prior to hybridization. Amplified cDNA (2000 ng per sample) with 10x Custom Blocker FWD (CTACACGACGCTCTTCCGATCT/3SpC3/) and 10x Custom Blocker REV (AAGCAGTGGTATCAACGCAGAG/3SpC3/) was used as an input for Twist custom probe overnight (18h) hybridization followed by capture of the hybridization mix with Twist streptavidin beads, washing of un-hybridized probes and on-beads post-capture PCR in order to increase the enriched cDNA mass (7 cycles). Twist-enriched full-length single cell cDNA was further used in Kinnex PCR with Kinnex primers, array formation and final Kinnex library generation following manufacture’s instructions from PacBio protocol ‘Procedure & checklist — Preparing Kinnex libraries using Kinnex single-cell RNA kit’. Kinnex libraries with Twist probe enrichment produced from sample A and sample B with the average size of 14 kb were quality checked, pooled and directed to DNA-polymerase complex preparation using Revio SPRQ binding chemistry and further sequenced on a single 25M Revio SMRT cell yielding 2.2 M HiFi reads/35 M segmented reads and 2.3 M HiFi reads/28 M segmented reads for sample A and sample B, respectively.

### Data Analysis

#### SR-WTA

A *CellRanger* ^[30]^ transcriptome reference was built from the human genome assembly GRCh38.p14 and the GENCODE v47 annotation using `cellranger mkref`. Pair-end scRNA-seq reads were processed with CellRanger v9.0.1 (`cellranger count`), which performs read alignment, barcode and UMI processing, cell calling, and PCR duplicate removal. The pipeline produced filtered gene–cell count matrices that were used for downstream analyses.

#### LR-WTA and LR-Twist

##### Preprocessing

The HiFi reads were deconcatenated into segmented reads (S-reads) based on segmentation adapters using SMRT Link. These S-reads were preprocessed following the Iso-Seq CLI workflow v4.3.0. This included primer removal using *lima* v2.12.0, followed by tagging, refinement, cell barcode correction and deduplication using *isoseq* v4.2.0.

##### Transcriptome alignment

We constructed the reference transcriptome from the GRCh38.p14 genome and the GENCODE v47 gene annotation using *gffread* v0.12.7^[31]^. Deduplicated reads were then aligned to this reference transcriptome with *minimap2* v2.28^[32]^ using the parameters `-ax map-hifi -N 100 --sam-hit-only --for-only`.

##### Genome alignment

We again used *minimap2* to align the deduplicated reads to the human reference genome (GRCh38.p14) with parameters `-ax splice:hq -uf --junc-bed`, using GENCODE v47 gene annotation BED as the junction reference.

##### Quantification

Transcriptome-aligned BAM files used quantified into transcript-by-cell count matrices using *oarfish* v0.7.0^[33]^, applying model coverage in single-cell mode without filtering and otherwise default settings. The resulting transcript-level count matrices were subsequently aggregated to generate gene-by-cell count matrices.

##### Downsampling analysis

We used *seqtk* v1.5^[34]^ to randomly subsample reads from SR-WTA FASTQ files using five independent seeds. The resulting downsampled FASTQ files were processed with *CellRanger*. In addition to the default cell-calling pipelines (*CellRanger* for SR-WTA and *isoseq* for LR-WTA), we applied the *emptyDropsCellRanger* function with default parameters from the *DropletUtils*^[20]^ package to the corresponding raw count matrices, including all barcodes.

##### Variant calling, filtering and annotation

We combined variants called by *Clair3* -*RNA* v0.2.2^[35]^ and *DeepVariant* v1.9.0^[36]^. Genome-aligned BAM files aggregated at the pseudobulk level were used as input for variant calling with *Clair3* - *RNA*^[35]^, applying the `hifi_mas_minimap2` model and argument `--tag_variant_using_readiportal` to annotate RNA-editing sites. The same BAM files were analyzed with *DeepVariant* using the MASSEQ model. Candidate variants were identified by filtering the raw calls from *Clair3* -*RNA* and *DeepVariant* independently. The following criteria were applied: (1) loci with fewer than five supporting reads at the pseudobulk level were excluded; (2) variants present in the dbSNP “common” subset were removed; (3) only variants marked PASS by the variant callers were retained, as non-PASS labels indicated low-quality calls or RNA-editing sites; and (4) only variants located within coding sequences (CDS) were considered. After filtering, VCF files were merged using the `merge` function in *bcftools* v1.21^[37]^, and read support at variant loci was assessed with `mpileup`. Finally, variants were annotated with *snpEff* v5.2^[38]^, and the predicted effect with the highest impact was reported.

##### Cell genotyping

For each candidate variant, we generated read pileups at the variant position and extracted cell bar-codes supporting the alternative allele(s) using R package *GenomicAlignments*^[39]^. A cell was classified as mutated for a given variant if at least one read carried the alternative allele, and as reference if only reads with the reference allele were observed. Because cells are diploid and coverage in scRNA-seq is limited, some cells classified as reference may in fact carry the mutation but lack reads covering the mutant allele^[3]^.

#### Cell typing for SR- and LR-WTA

Doublet cells were first identified and removed using *scDblFinder* ^[40]^, incorporating sample information since cells were captured separately for each sample. We then applied quality control and filtering with the *scater* ^[41]^ R package. Genes that were not detected in any cell were first removed. Cells with total counts, number of detected features, or percentage of mitochondrial genes exceeding 2.5 median absolute deviations from the median were then filtered out. Finally, we only kept genes with a count greater than 1 in at least 20 cells for downstream analysis.

Next, we applied *scran*’s *modelGeneVar* ^[42]^ function to identify the 3,000 most highly variable genes (HVGs) using default parameters. These HVGs were then used for dimensionality reduction with PCA and UMAP, based on the first 20 principal components. Cell annotation was performed using *CellTypist* ^[22][23]^ with the *Fetal_Human_AdrenalGlands* model^[24]^. To further resolve cell subtypes, we applied the *Immune_All_High* model to re-annotate cells initially classified as myeloid. *CellTypist* assigns a confidence score to each annotation, defined as the probability of the predicted cell type from the logistic regression model.

## Availability of data and materials

The *Snakemake* workflow and the code used to generate the figures are available on [GitHub](https://github.com/Jiayi-Wang-Joey/hybrid-sequencing). The expression count tables are available at [Zenodo](https://zenodo.org/records/17494517). The raw sequencing data have been deposited in the European Genome-phenome Archive (EGA) under accession EGAS50000001537.

## Authors’ contributions

F.B. provided access to the adrenal samples, and M.C.M. processed them. Q.Z. and A.B.N. contributed to conception of the experiment. A.B.N., M.C.M. and Q.Z. coordinated the sequencing experiment. J.W. wrote the Snakemake workflow and performed all data analyses with input from M.D.R. J.W. wrote the original manuscript, with contributions from M.C.M. and A.B.N. on experimental methods, and with revisions and edits from M.D.R. M.D.R., D.P. and F.B. acquired funding and co-supervised the work. All authors reviewed, edited, and approved the final manuscript.

## Acknowledgments

We thank the members of the Robinson Lab and the Electrophysiology Facility at the University of Zürich for their valuable feedback on data analysis, especially David Wissel for long-read data processing and Dr. Izaskun Mallona for advice on variant calling. We acknowledge Dilmurodjon Eshmuminov, Diana Vetter, and Umberto Maccio at the University Hospital Zürich for their support during clinical inspections. We also acknowledge Tamara Carrasco Oltra for valuable support with single-cell experiments at the Functional Genomics Center Zürich (FGCZ, ETHZ and UZH). Finally, we thank the sequencing facilities for generating the PacBio sequencing data (Next Generation Sequencing Platform, University of Bern, Switzerland) and the Illumina sequencing data (FGCZ).

## Ethics approval and consent to participate

This study was approved by the Cantonal Ethics Committee of Zürich (Kantonale Ethikkommission Zürich), Switzerland (BASEC No. 2017–00771).

## Consent for publication

Not applicable; no identifiable individual data are included.

