## Supplementary Material for "Hybrid untargeted and targeted RNA sequencing facilitates genotype-phenotype associations at single-cell resolution"

[1]*Department of Molecular Life Sciences, University of Zurich, Zurich, Switzerland*
[2]*SIB Swiss Institute of Bioinformatics, University of Zurich, Switzerland*
[3]*Instituto de Fisiología, Facultad de Ciencias, Universidad de Valparaíso, Valparaíso, Chile*
[4]*Centro Interdisciplinario de Neurociencia de Valparaíso, Facultad de Ciencias, Universidad de Valparaíso, Valparaíso, Chile*
[5]*Functional Genomics Center Zurich, ETH Zurich and University of Zurich, Zurich, Switzerland*
[6]*Department of Endocrinology, Diabetology and Clinical Nutrition, University Hospital Zurich (USZ), 8091 Zurich and University of Zurich (UZH), 8006 Zurich, Switzerland*
[7]*Medizinische Klinik und Poliklinik IV, Klinikum der Universität, Ludwig-Maximilians-Universität, D-80336 Munich, Germany*
[8]*The LOOP Zurich - Medical Research Center, Zurich, Switzerland*
[9]*Electrophysiology Facility, University of Zurich, Switzerland*

### 1 Supplementary tables

#### 1.1 Data quality

| Workflow | CellRanger | isoseq |
|---|---|---|
| Primer removal | ✓ | ✓ |
| Cell barcode correction | ✓* | ✓† |
| CB whitelist | ✓ | ✓ |
| UMI correction | ✓ | ✗ |
| Deduplication | ✓* | ✓† |
| Cell calling | ✓* | ✓† |
| Reselect sub-threshold cells | ✓ | ✗ |

**Table S1:** Summary of processing steps with CellRanger for SR-WTA and isoseq for LR-WTA/Twist. * after mapping; † before mapping.

| **Metric** | **Sample A** | **Sample B** |
|---|---|---|
| Number of Reads | 256,751,486 | 228,842,856 |
| Estimated Number of Cells | 10,174 | 5,687 |
| Mean Reads per Cell | 25,236 | 40,240 |
| Median Genes per Cell | 1,154 | 1,966 |
| Valid Barcodes | 89.2% | 90.4% |
| Valid UMIs | 99.9% | 99.9% |
| Sequencing Saturation | 31.1% | 41.7% |
| Q30 Bases in Barcode | 96.5% | 96.0% |
| Q30 Bases in RNA Read | 95.2% | 94.2% |
| Q30 Bases in UMI | 97.4% | 97.4% |
| Reads Mapped to Genome | 95.1% | 94.9% |
| Reads Mapped Confidently to Genome | 68.2% | 67.1% |
| Reads Mapped Confidently to Intergenic Regions | 1.0% | 0.8% |
| Reads Mapped Confidently to Intronic Regions | 18.7% | 15.0% |
| Reads Mapped Confidently to Exonic Regions | 48.5% | 51.3% |
| Reads Mapped Confidently to Transcriptome | 60.6% | 60.7% |
| Reads Mapped Antisense to Gene | 4.6% | 3.6% |
| Fraction Reads in Cells | 60.4% | 66.8% |
| Total Genes Detected | 44,778 | 40,835 |
| Median UMI Counts per Cell | 2,191 | 4,642 |

**Table S2:** CellRanger summary metrics for Sample A and Sample B for SR-WTA.

| Type | Sample | LR-WTA | LR-Twist |
|---|---|---|---|
| raw | sampleA | 85,182,821 | 35,472,753 |
| raw | sampleB | 78,196,246 | 28,235,749 |
| deduplicated | sampleA | 74,820,537 | 7,217,104 |
| deduplicated | sampleB | 67,218,222 | 5,538,856 |
| cell_calling | sampleA | 26,999,791 | 3,613,091 |
| cell_calling | sampleB | 28,196,044 | 3,187,328 |
| mapped_genome | sampleA | 26,862,528 | 3,591,677 |
| mapped_genome | sampleB | 28,180,994 | 3,185,666 |
| mapped_transcriptome | sampleA | 20,295,857 | 2,541,682 |
| mapped_transcriptome | sampleB | 22,538,341 | 2,490,549 |
| counts | sampleA | 19,638,599 | 2,541,073 |
| counts | sampleB | 21,785,336 | 2,489,980 |

**Table S3:** Number of reads at each processing step in the IsoSeq workflow for LR-WTA/Twist samples, following the standard *isoseq* analysis pipeline.

#### 1.2 Strand bias

*CYP11B2* lies on the negative genomic strand. In a strand-specific library such as MAS-Seq, reads from *CYP11B2* transcripts are therefore expected to align predominantly to the reverse strand, and the observed strand asymmetry at this locus is consistent with that orientation. Strand specificity fixes the *global* orientation, but strand-bias tests (Fisher exact test) ask whether REF and ALT share the same +/− distribution among the reads that do occur. Even in stranded data, a small fraction of opposite-strand alignments is expected (e.g., rare antisense, capture effects); if ALT is disproportionately enriched in that minority strand while REF is not, this suggests an assay/mapping artifact rather than biology. To evaluate strand bias at *CYP11B2* (chr8:142,912,613 G>A) from the IGV observations, we formed a $2 \times 2$ table of forward/reverse counts for REF and ALT and applied Fisher's exact test. The Fisher strand score (FS) was computed as:

$$\mathrm{FS} = -10 \times \log_{10}(p)$$

where $p$ is the two-sided $p$-value from Fisher's exact test.
For the LR-WTA dataset, the counts were:

| | Forward | Reverse |
|---|---|---|
| REF (G) | 1 | 15 |
| ALT (A) | 1 | 5 |

yielding $p = 0.48$ and $\mathrm{FS} = -10 \times \log_{10}(0.48) = 3.18$.
For the LR-Twist dataset, the counts were:

| | Forward | Reverse |
|---|---|---|
| REF (G) | 8 | 300 |
| ALT (A) | 5 | 43 |

yielding $p = 0.02$ and $\mathrm{FS} = -10 \times \log_{10}(0.02) = 16.95$.

#### 2 Supplementary figures

##### 2.1 Data analysis workflow

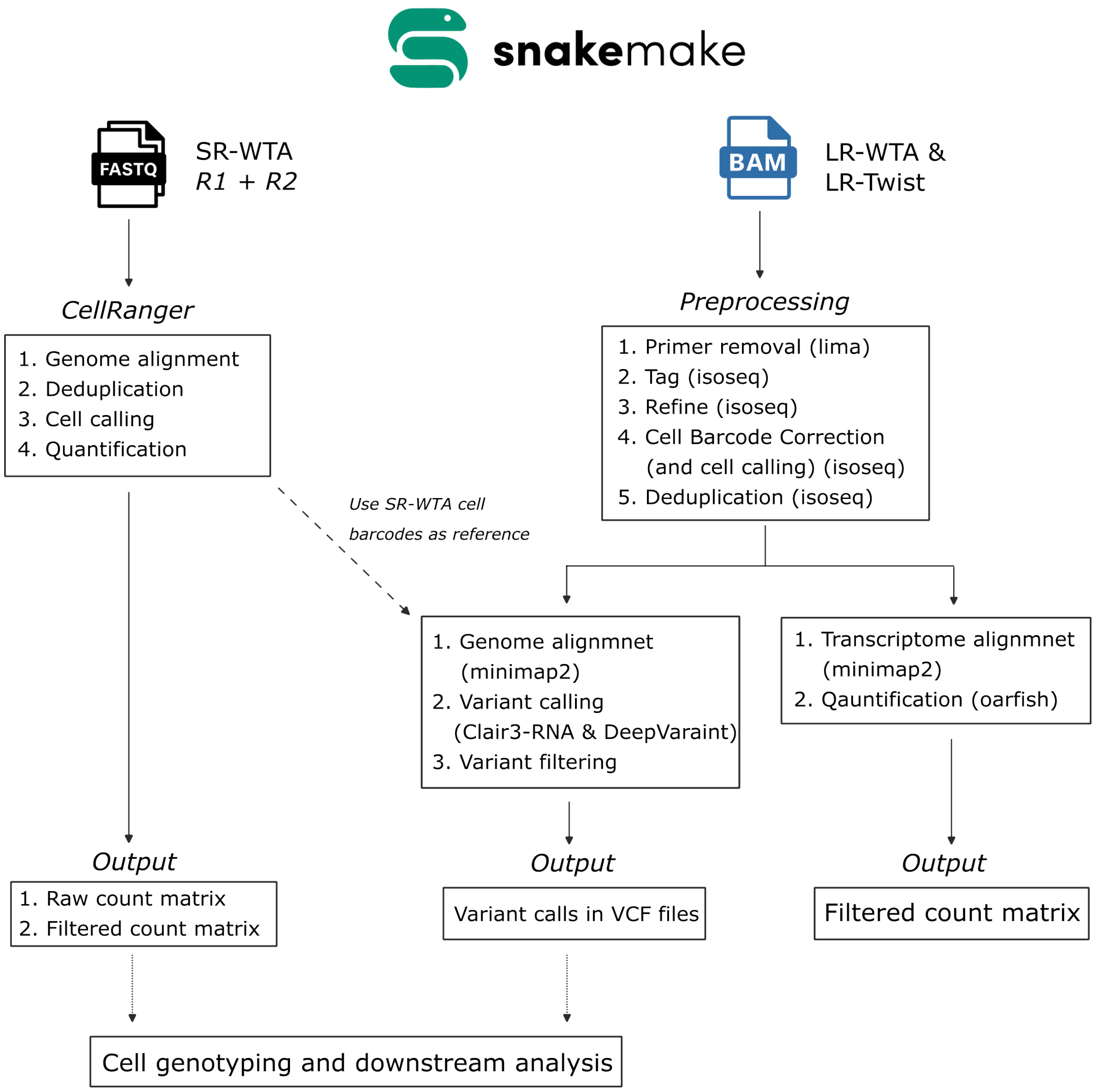

**Fig. S1: Schematic Snakemake workflow of data analysis.**

##### 2.2 Data quality

We visualized the number of reads retained at each processing step for SR-WTA, LR-WTA, and LR-Twist (Fig. S2). Both *CellRanger* and *isoseq* use a knee-finding method by default to determine the cell barcode threshold. However, for LR-Twist datasets, this approach underestimated the true number of cells because the barcode rank plots displayed a gradual curve rather than a sharp inflection point. To address this, we applied a percentile-based thresholding method using the 95th percentile for the long-read datasets. With this 95% cutoff, the number of detected cells in LR-WTA closely matched that obtained using the knee-finding method. For SR-WTA, since CellRanger does not produce intermediate BAM files, the number of reads at each step was estimated from the metrics reported in Table S2.

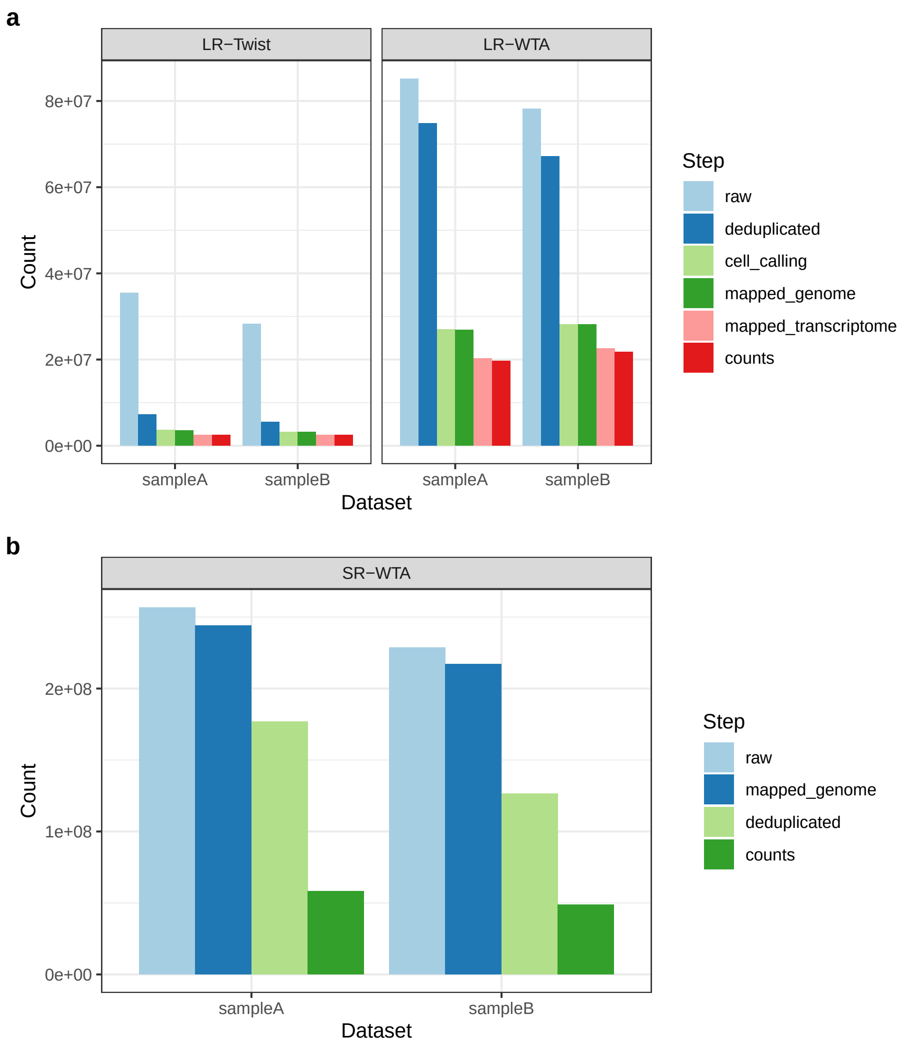

**Fig. S2: The number of reads at each data processing step.**

#### 2.3 SR-WTA vs. LR-WTA

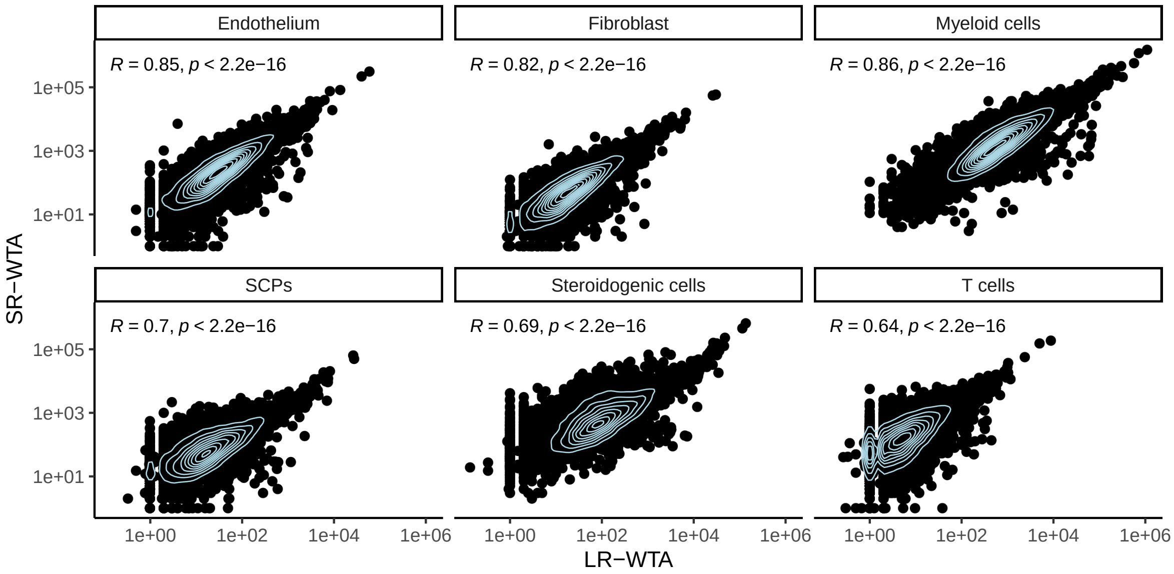

**Fig. S3: Spearman's correlation on UMI counts per gene between SR- and LR-WTA, stratified by cell types.**

**Fig. S4: Comparison of downsampled SR-WTA and LR-WTA using default cell-calling methods.** (**a**) Cell barcode rank plots, where the x-axis shows barcodes ranked in descending order and the y-axis represents the total UMI counts per barcode. Points are colored according to whether they were called as cells by the respective default pipelines, i.e., CellRanger for SR-WTA and isoseq for LR-WTA. (**b**) Venn diagrams illustrating the overlap between downsampled SR-WTA and LR-WTA cells, with the first row corresponding to sampleA and the second to sampleB. (**c**) UMAP projections of cells from sampleA and sampleB, colored by cell types annotated using CellTypist. (**d**) Bar plots showing cell type proportions in LR-WTA and downsampled SR-WTA. Panels **c** and **d** share the same color legend.

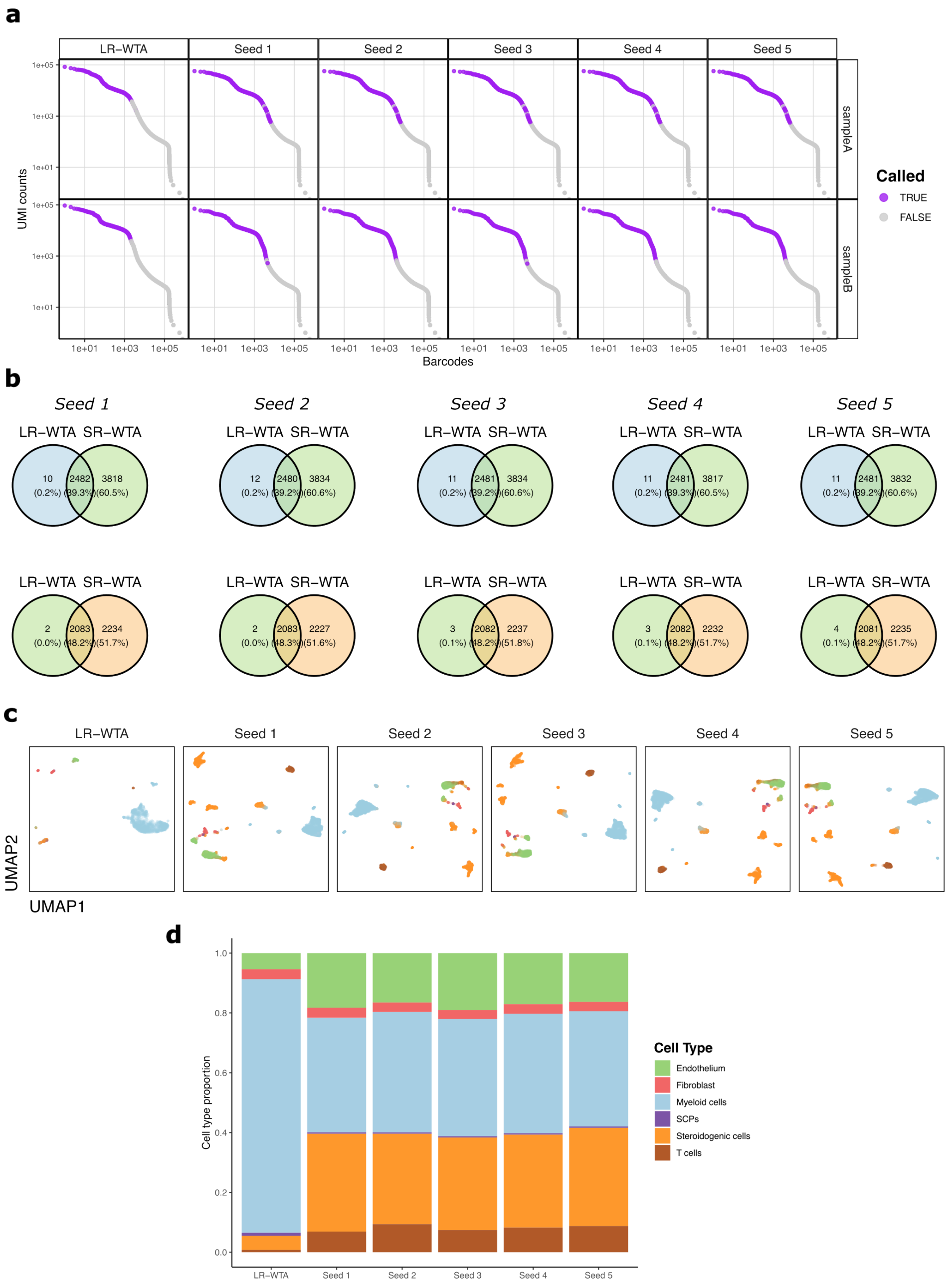

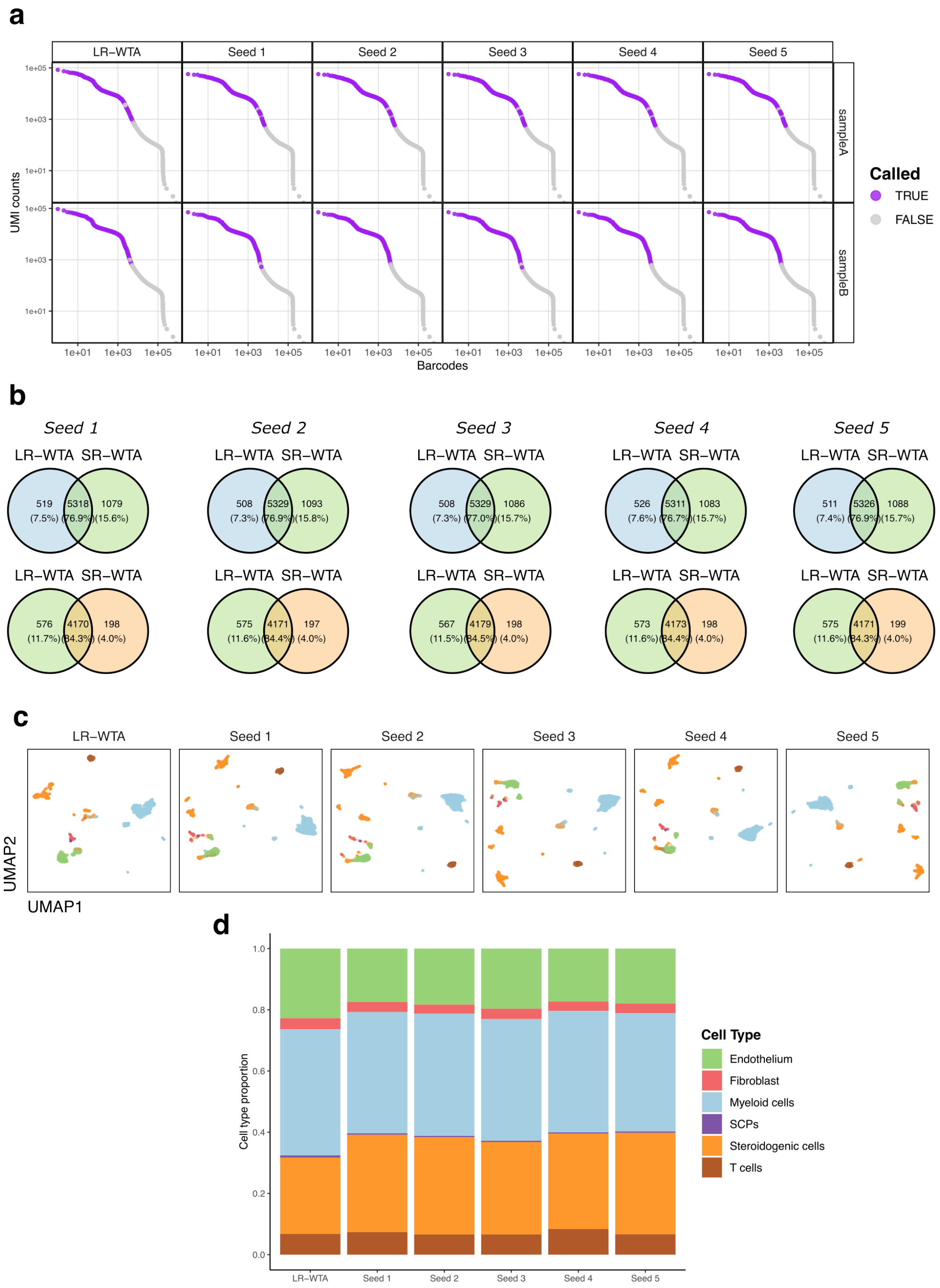

**Fig. S5: Comparison of downsampled SR-WTA and LR-WTA using the same cell calling method.** Same as Fig.S4, but the same cell-calling method (EmptyDrops) was applied to the raw count matrices.

#### 2.4 LR-WTA vs. LR-Twist

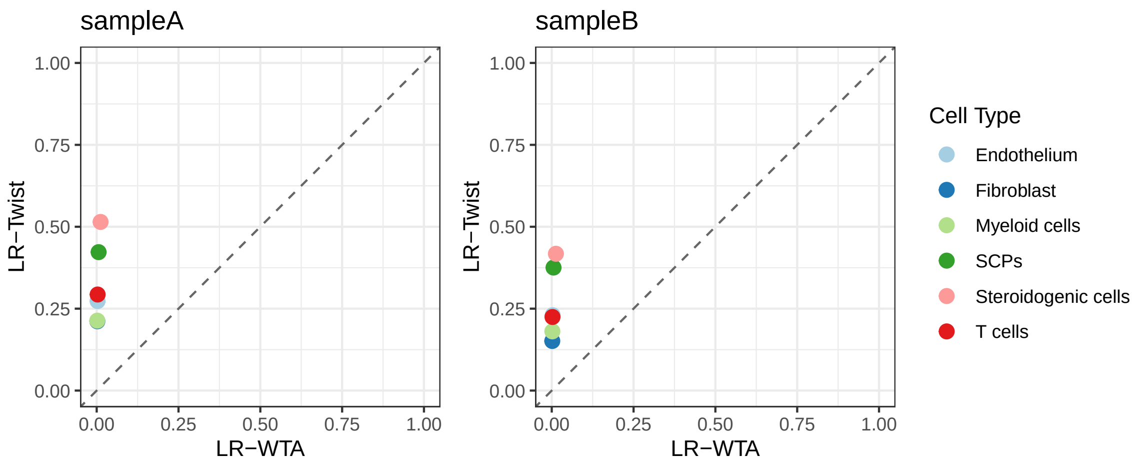

**Fig. S6: Proportion of targeted reads in each cell type in LR-WTA and LR-Twist, colored by cell types.**

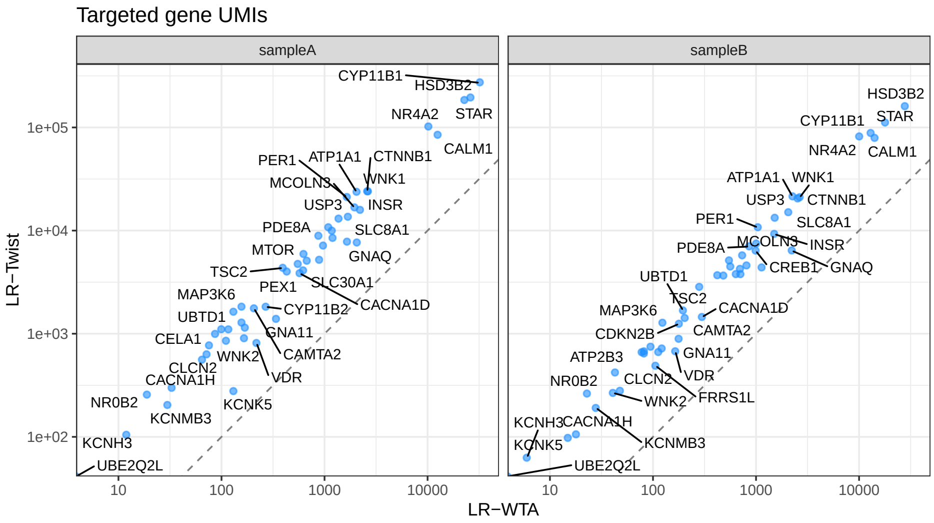

**Fig. S7: Twist capture efficiency on targeted genes.** Scatter plot comparing summed gene-level UMI counts across all cells between LR-WTA and LR-Twist (on log10 scale), only for targeted genes.

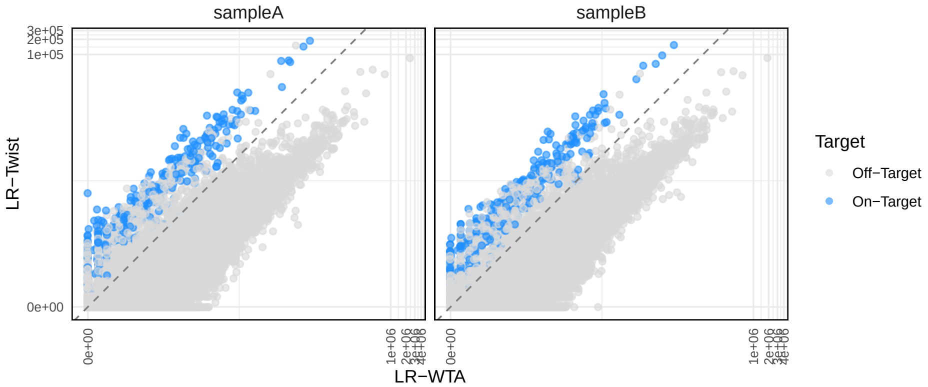

**Fig. S8: Twist capture efficiency on targeted transcript.** Scatter plot comparing summed transcript-level UMI counts across all cells between LR-WTA and LR-Twist (on log10 scale).

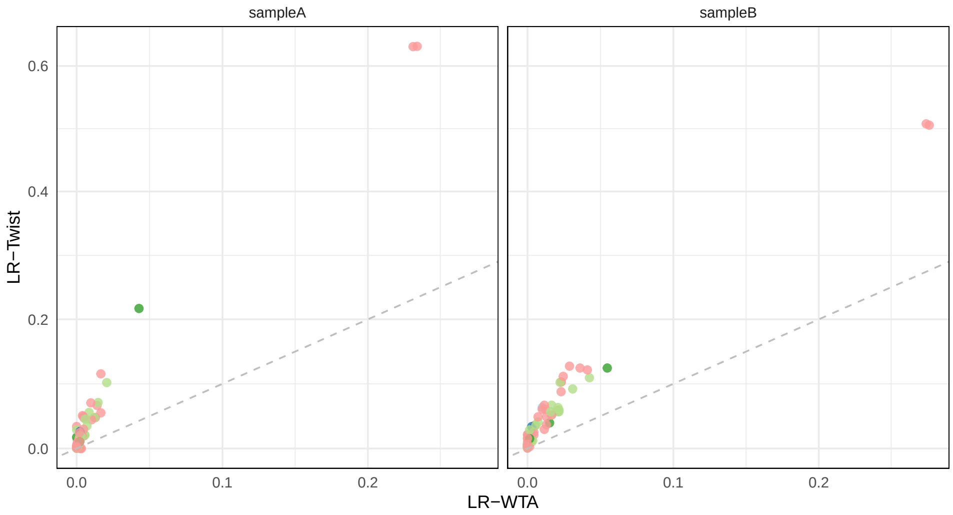

**Fig. S9: Proportion of steroidogenic cells genotyped in LR-WTA and LR-Twist.** The total number of steroidogenic cells is the same in LR-WTA and LR-Twist, since both are based on the cells identified in SR-WTA.

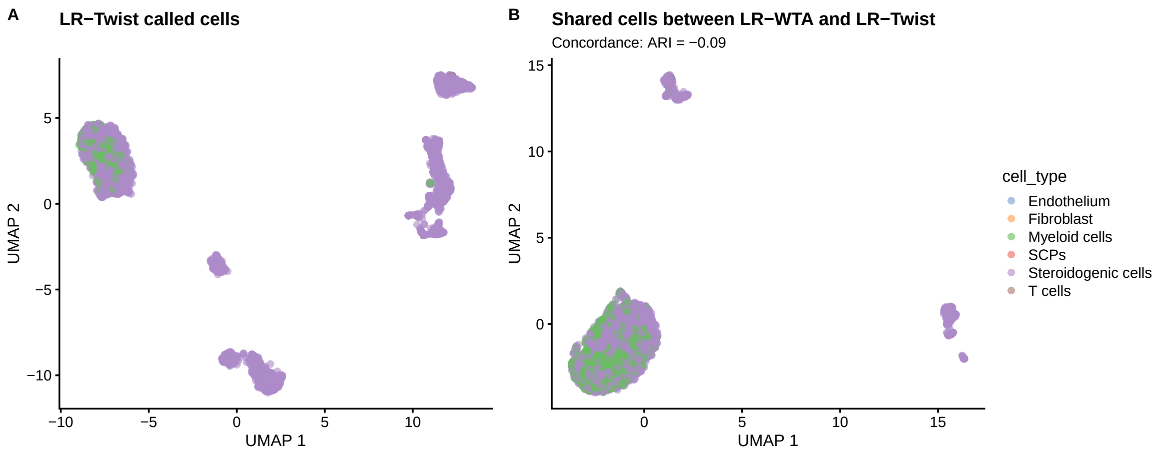

**Fig. S10: UMAP projection of cells from samples A and B for LR-Twist.** (**a**) UMAP of called cells from LR-Twist. (**b**) UMAP of cells shared between LR-WTA and LR-Twist, annotated with the adjusted Rand index (ARI) computed from the cell type labels of LR-Twist and LR-WTA. Low-dimensional embeddings were generated using the top 3,000 highly variable genes (HVGs), and cells are colored by cell types assigned by `CellTypist`.

Here, we used one gene as an example to illustrate the inefficiency of variant calling from SR-WTA. In Integrative Genomics Viewer (IGV), reads from SR-WTA are concentrated at the 3' end and only a few are mapped to the coding regions, in contrast to LR-WTA and LR-Twist, which show broader coverage across the gene body.

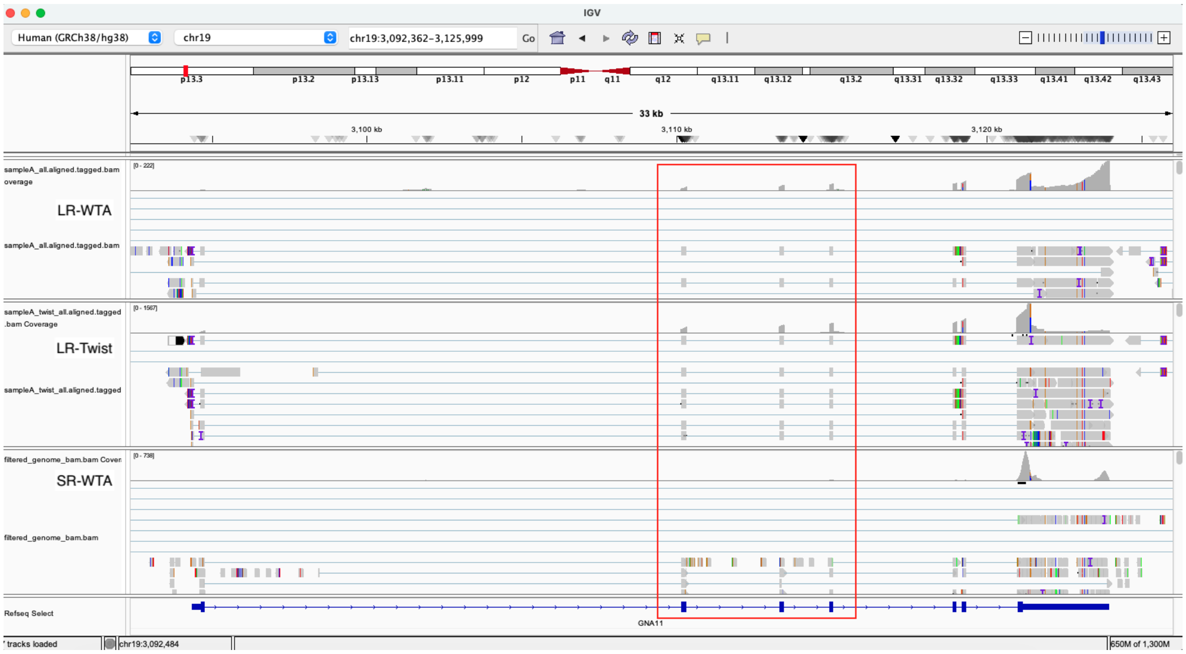

**Fig. S11: IGV screenshots of reads mapped to *GNA11* from SR-WTA, LR-WTA and LR-Twist.**
